# Different impacts of naturally occurring variants of the globally disseminated carbapenemase-encoding plasmid pKpQIL on the biology of host strains

**DOI:** 10.1101/495986

**Authors:** Howard T. H. Saw, Mark A. Webber, Neil Woodford, Laura J. V. Piddock

**Affiliations:** Antimicrobials Research Group, Institute of Microbiology & Infection, College of Medical & Dental Sciences, University of Birmingham, Edgbaston, United Kingdom.; NIS Laboratories, National Infection Service, Public Health England, London, United Kingdom.

**Keywords:** Plasmid, KPC, pKpQIL, fitness, carbapenem

## Abstract

*Klebsiella-associated* plasmid pKpQIL and its variant have been isolated globally. Our study aimed to determine whether a naturally occurring variant has altered host range and impacts on the fitness of different bacterial host strains. The plasmids pKpQIL-UK and pKpQIL-D2 were transferred from the original clinical isolate host strains of *Klebsiella pneumoniae* into *Escherichia coli, Salmonella* Typhimurium, *Enterobacter cloacae* and *Serratia marcescens* strains by filter-mating and conjugation frequencies determined and compared. The fitness of the resulting transconjugants was assessed by determining growth kinetics, ability to form a biofilm and persistence of the plasmids in each host was also measured. Transfer of either plasmid into *Salmonella* and S. *marcescens* was similar. However, pKpQIL-UK transferred into *E. coli* at a higher rate than did pKpQIL-D2; the reverse was found for *E. cloacae*. Both plasmids were rapidly lost from the *E. coli* population. Plasmid pKpQIL-UK, but not -D2, was able to persist in *Salmonella*. Although pKpQIL-UK imposed a greater fitness cost (inferred from an increased generation time) than -D2 on *E. cloacae*, it was able to persist as well as pKpQIL-D2 in this host. The pKpQIL-D2 plasmid did not confer any fitness benefit on any of the hosts under the conditions tested. Variants of the globally important pKpQIL plasmid have arisen in patients due to recombination. The impacts of the pKpQIL-UK plasmid and the -D2 variant in various Enterobacteriaceae are host-dependent. Continuing evolution of pKpQIL may alter its host range in the future.

## INTRODUCTION

The World Health Organisation (WHO) has placed carbapenem- and 3^rd^-generation cephalosporin-resistant Enterobacteriaceae *(Klebsiella pneumoniae, Escherichia coli, Enterobacter* spp. and *Serratia* spp.) in the ‘Critical Priority Pathogens List’, emphasising the urgent need to find drugs active against these bacteria (1). In 2018, the WHO’s latest antibiotic resistance surveillance report continued to show high rates of antibiotic resistance across various high- and low-income countries, with *K. pneumoniae* as the most common species of carbapenem-resistant Enterobacteriaceae (2).

Since they were first identified two decades ago (3), *K. pneumoniae* carbapenemases (KPC) have become the most widespread carbapenemases in several regions across the world (4-6). The spread of the KPC enzymes has been largely associated with expansion of the hyperendemic clone, *K. pneumoniae* ST258 and its single locus variants (7). Worryingly, KPC enzymes are increasingly described in various Enterobacteriaceae members, including *E. coli, Enterobacter cloacae, Enterobacter aerogenes* and *Serratia marcescens* (8, 9). In addition to β-lactam antibiotics, such carbapenemase producers are often resistant to other unrelated groups of antibiotics, including colistin, fluoroquinolones and aminoglycosides, limiting the availability of treatment options (10, 11). In *K. pneumoniae*, the successful dissemination of KPC carbapenemase has often been attributed to the spread of the *K. pneumoniae-adapted* pKpQIL plasmid and very closely related ‘pKpQIL-like plasmids’ which vary from the original by only a few single nucleotide polymorphisms (SNPs) (12). pKpQIL-like plasmids have been connected to hospital outbreaks of both ST258 and non-ST258 *K. pneumoniae*, as well as various other Gram-negative bacteria, including the pandemic, human-adapted and extra-intestinal pathogenic *E. coli* (ExPEC) lineage, ST131 (12-14).

Initially discovered in Israel (15), the origin of the 114 kb IncFII plasmid pKpQIL has been traced back to the United States (12). Apart from conferring carbapenem resistance, it has also been suggested that pKpQIL influences the resistome of its bacterial host due to the transposition of insertion elements from the plasmid into chromosomal genes (e.g *mgrB)* leading to additional resistance (10). The *bla*_KPC_ gene is contained within a *Tn4401a* transposon, which is itself independently mobile and can become integrated into a secondary transposon (9).

As with many other regions around the world (12, 16), KPC-producing *K. pneumoniae* have been isolated in the UK and pKpQIL-UK (which differs from the original Israel pKpQIL by a few base substitutions) has been identified in major hospital outbreaks of carbapenem-resistant *K. pneumoniae* in the country (17, 18). In one UK region, both pKpQIL-UK and a variant where a 19.5 kb region in the former has been substituted with a 17.6 kb fragment have been found to be in circulation in isolates from patients (17). This variant, named ‘pKpQIL-D2’ still carries the *bla*_KPC_ gene and was found in multiple species of Enterobacteriaceae (14, 17). The substituted region gained by the -D2 variant contains 19 putative open reading frames including putative partitioning proteins *(parA* and *parB)*, transposases, a resolvase and replication initiator protein as well as several predicted proteins with no known function (17).

Other rearrangements of the pKpQIL backbone have also been reported elsewhere (12, 13, 16), indicating the dynamic genetic nature of this plasmid and suggesting that these variants may be the result of selection for adaptation to the host in a niche environment.

Previously, our team has shown that both the pKpQIL-UK and -D2 are well adapted to their *Klebsiella* hosts and transcriptional changes were effected to ameliorate (if any) possible fitness cost from plasmid carriage in ST258 strains of *K. pneumoniae* (19). Given the identification of the -D2 variant in various species, we hypothesised that the 17.6 kb variable region within pKpQIL-D2 may provide an adaptive fitness advantage when compared with the parent plasmid, which facilitated a broader host range and increased dissemination potential than the original pKpQIL in the UK. We tested this hypothesis by measuring: (i) the transfer of the two plasmids into new Enterobacteriaceae hosts; (ii) the generation times of new host +/- plasmid; (iii) the ability of new host +/- plasmid to form biofilm; (iv) the minimum inhibitory concentrations (MIC) of the plasmid-carrying Enterobacteriaceae; and (v) the persistence of the plasmids in their new hosts.

## RESULTS

### Conjugation frequencies of the plasmids varies across Enterobacteriaceae

The ability of the plasmids (pKpQIL-UK and -D2) to transfer from their respective original host clinical isolates was investigated. There were different conjugation frequencies from the original *K. pneumoniae* isolates into the different species of Enterobacteriaceae tested. There was no significant difference detected in the frequency of transfer of the plasmids into S. Typhimurium (Table 1). However, the pKpQIL-UK plasmid transferred into *E. coli* at a frequency 33-fold higher than pKpQIL-D2. Interestingly, the opposite was observed with *E. cloacae* where the pKpQIL-D2 transferred at a higher frequency (11-fold) into this species than the pKpQIL-UK. The pKpQIL-D2 also transferred into *S. marcescens* at a higher frequency (7-fold) than the -UK plasmid although this difference was not statistically significant.

**Table 1.**
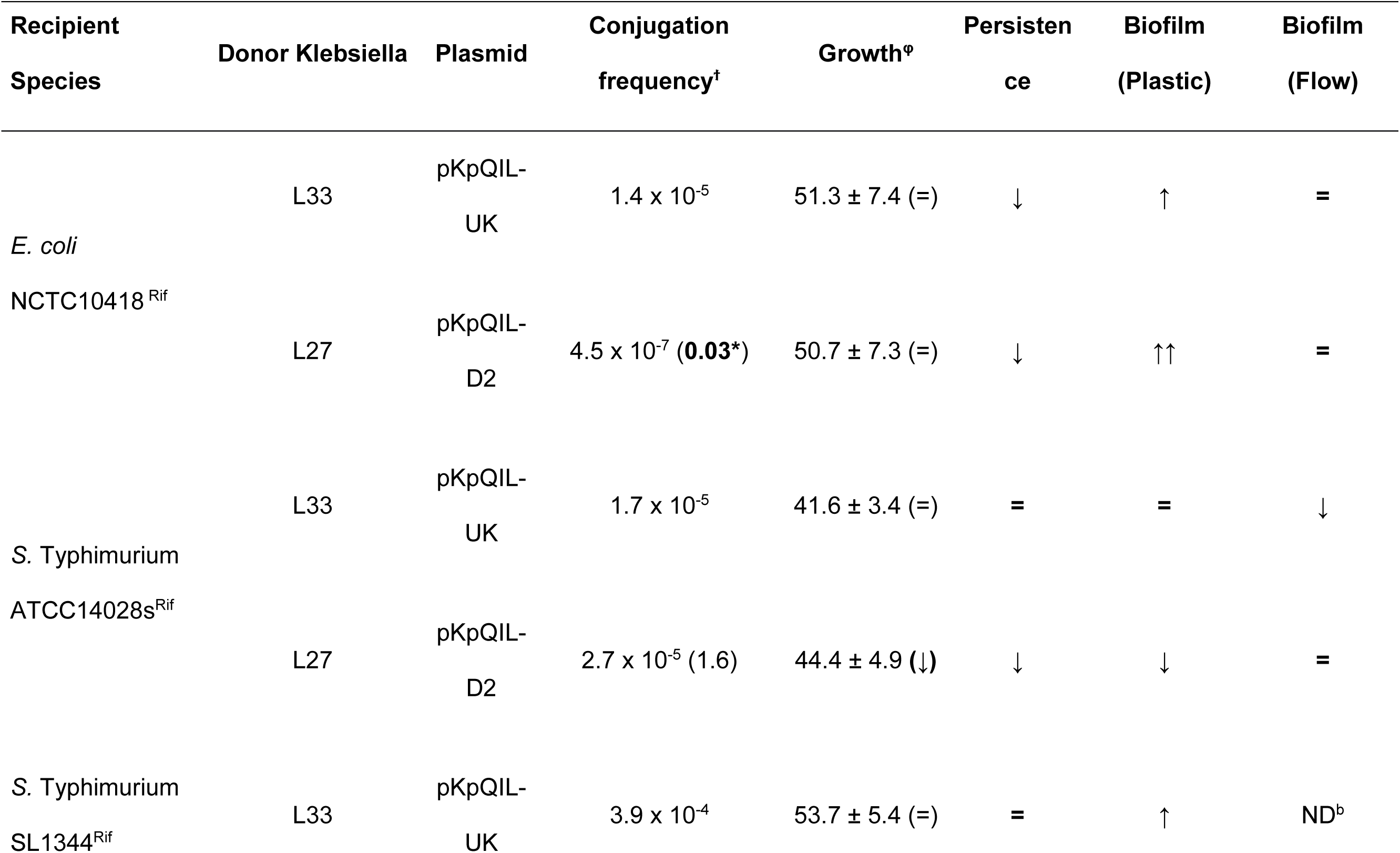
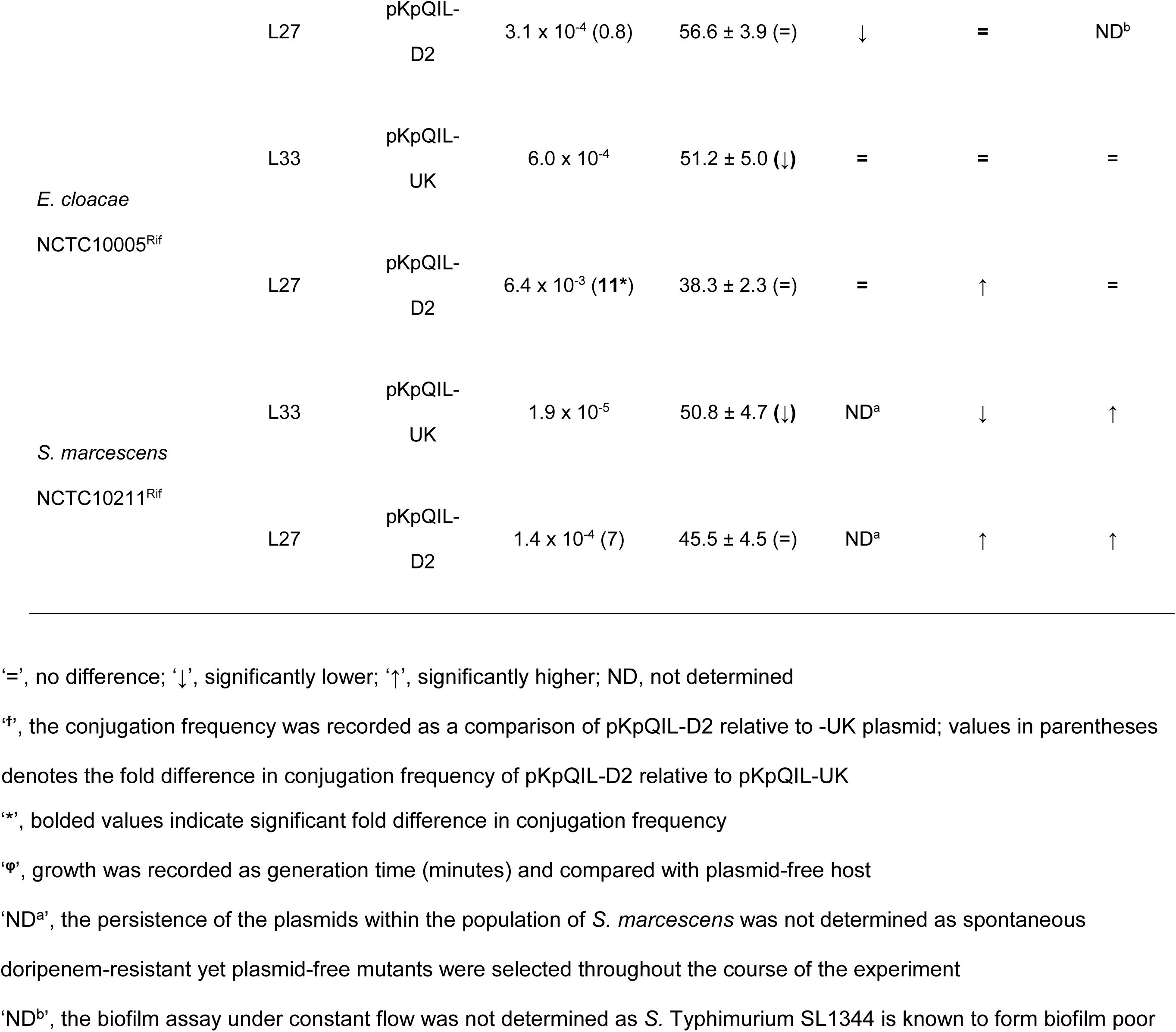
Summary of the impact of pKpQIL-UK and pKpQIL-D2 upon the plasmid-carrying Enterobacteriaceae.

### Plasmid carriage did not confer an altered generation time on host strains

It was hypothesised that the pKpQIL-D2 plasmid may impact fitness of host strains differentially to pKpQIL-UK. Hence, the generation time of the various plasmid-carrying Enterobacteriaceae was compared with the isogenic plasmid-free strain. In *E. coli* and S. Typhimurium SL1344, no difference in generation time was observed between the plasmid-carrying and plasmid-free bacteria. Interestingly, the S. Typhimurium ATCC14028s carrying the pKpQIL-D2 plasmid had a longer generation time than the plasmid-free isogenic strain. In contrast, both *E. cloacae* and S. *marcescens* carrying the pKpQIL-UK plasmid had longer generation times than the plasmid-free strains whereas these strains were not significantly affected by carriage of the -D2 variant (Table 1).

### Plasmids pKpQIL-UK and pKpQIL-D2 persisted differently in different host strains

For a plasmid to be successful within a bacterial population, it must be stably maintained within its host. Therefore, the ability of each plasmid to persist without any antibiotic pressure was investigated over a period of 20 days (ca. 140 generations) (Figure 1 and Table 1). Both plasmids were rapidly lost from the *E. coli* population with approximately 75% of the population retaining the plasmids after 5 days. At the end of the 20-day period, no *E. coli* carried the plasmids. In S. Typhimurium, pKpQIL-UK persisted successfully without detectable loss at the end of the experiment. However, pKpQIL-D2 did not persist within the *Salmonella* population. *E. cloacae* retained both plasmids after ca. 140 generations. The persistence of the plasmids within *S. marcescens* was not determined as spontaneous doripenem-resistant mutants were indirectly selected during the course of the experiment.

**Figure 1.**
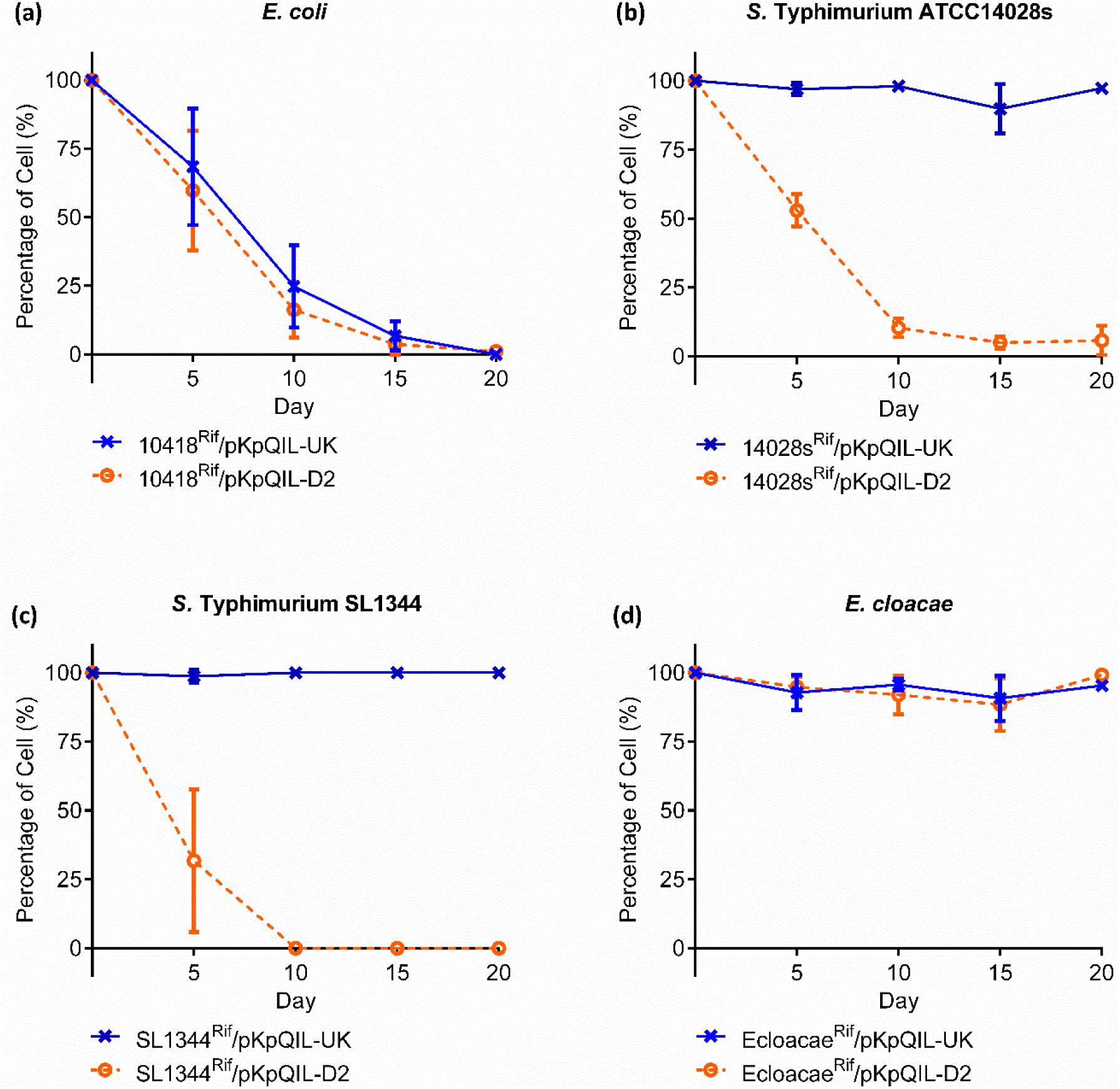
The percentage of pKpQIL-UK vs -D2 carrying cells in growth over a period of 20 days without antibiotic selection. The persistence of the plasmids pKpQIL-UK (Blue cross) and pKpQIL-D2 (Orange open circle) in various Enterobacteriaceae species in LB broth without antibiotic selection over a 20-day period was investigated to determine the stability of the plasmids in their respective hosts. The percentages of cells which retain the plasmids were recorded as mean ± standard deviation of three independent experiments.

### Plasmid-carrying Enterobacteriaceae form biofilm differently

As many bacteria exist as a biofilm rather than planktonic state in the natural environment, we investigated the contribution of the plasmids to the host bacterium’s ability to form a biofilm. Although both plasmids increased the ability of *E. coli* to form a biofilm on plastic, the pKpQIL-D2-carrying *E. coli* formed significantly more biofilm (Figure 2 and Table 1). Interestingly, the opposite was observed with *S*. Typhimurium. We studied two strains of *S*. Typhimurium which have different inherent capacities to form biofilms; 14028s considered a proficient biofilm forming strain and SL1344, a weak biofilm forming strain (20). pKpQIL-D2 reduced the ability of 14028s to form a biofilm, while pKpQIL-UK increased the ability of SL1344 to form a biofilm (although this improved level was still lower than 14028s). Compared with the plasmid-free host bacteria, pKpQIL-D2-carrying *E. cloacae* and S. *marcescens* formed more biofilm in the microtitre tray assay. In contrast, pKpQIL-UK did not change the ability of *E. cloacae* to form a biofilm but significantly reduced the amount of biofilm formed by *S. marcescens*.

**Figure 2.**
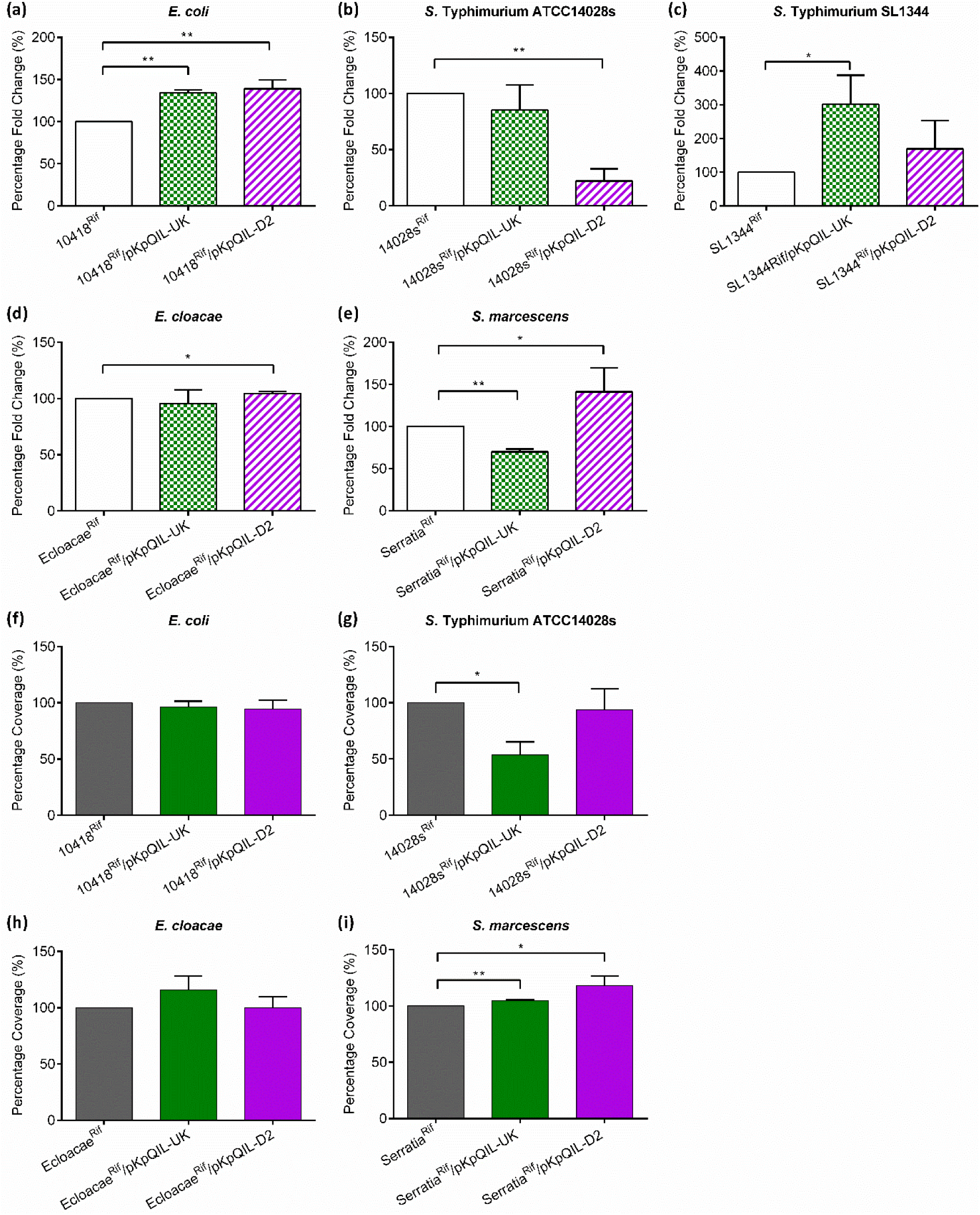
Biofilm formation for the various plasmid carrying Enterobacteriaceae strain. The impact of the plasmids pKpQIL-UK (Chequered) and pKpQIL-D2 (Striped) on its hosts’ ability to form biofilm was investigated using both the microtitre tray ‘crystal violet’ method and by determining the ability to cover flow cells in a microfluid platform. Panels a-e show crystal violet data and indicate percentage fold change compared to plasmid-free parent strain. Data show mean values ± standard deviation of three independent experiments. Panels f-I show percentage area of coverage by biofilm in microfluidic channels after 48 hours under constant flow of LB broth (without sodium chloride). The coverage of biofilm formed by the various pKpQIL-UK (Green) and pKpQIL-D2 (Magenta) carrying strains on the surface of the microfluidic channel was estimated using ImageJ software. The percentage coverage relative to the parental strain was recorded as mean ± standard deviation of three independent experiments. For both data sets student’s *t-*test was used to analyse significant changes and shown with asterisk (*) p<0.05 and (**) p<0.001.

Under constant flow, pKpQIL-UK decreased the amount of biofilm formed by *S*. Typhimurium ATCC14028s but increased the amount of biofilm formed by *S. marcescens* observed in the microfluidic channel (Figure 2). The pKpQIL-D2 plasmid also increased the ability of *S. marcescens* to form a biofilm under constant flow but had no significant effect upon biofilm formation by the other Enterobacteriaceae hosts. The biofilm formation of *S*. Typhimurium SL1344 was not assessed under flow as this strain did not efficiently colonise the flow cells (20).

### Susceptibility to antibiotics

With the exception of *S. marcescens*, MICs were the same irrespective of whether pKpQIL or pKpQIL-D2 were carried by the hosts bacterial strains (Table S1). In *S. marcescens*, pKpQIL-UK conferred significantly higher carbapenem resistance (4- to 8-fold increases in MICs of ertapenem, imipenem, meropenem, doripenem and biapenem). Although it has been previously reported that down-regulation of porins can result in carbapenem resistance in *S. marcescens*, we found no difference in the amount of porins expressed by the plasmid-carrying strains (data not shown).

## DISCUSSION

Since the discovery of the pKpQIL in Israel (15), this plasmid and its variants have been isolated in bacteria from various continents including North America, Europe and Asia (6, 21-23). Although the *bla*_KPC_ carbapenemase can be found in various Enterobacteriaceae (22), it is predominantly associated with *K. pneumoniae* and the pKpQIL plasmid (7, 24). This plasmid has a high propensity for genetic rearrangements (12, 13, 16), but the benefit of such genetic plasticity and consequences of evolution of the plasmid in relation to different hosts remain unclear. Here, we investigated whether the pKpQIL-D2 plasmid confers a different fitness impact to that of the more widely observed pKpQIL-UK plasmid.

Bacterial fitness is generally perceived as the adaptation of the bacterium towards its surrounding conditions, in order for its genetic traits to be maintained in a population either through inheritance by its progeny or horizontal gene transfer into different hosts (25, 26). It is typically accepted that plasmid carriage impacts the bacterial host negatively through the costs incurred on the replication, transcription and translation machineries, leading to the preferential selection of plasmid-free hosts in the population (27-30). In this study, the fitness of the plasmid-bearing strains was investigated in various ways, such as growth kinetics, biofilm formation, conjugation frequency and antibiotic susceptibility.

The growth of strains carrying pKpQIL-UK and -D2 were indistinguishable from those of plasmid-free strains for *S*. Typhimurium SL1344 and *E. coli* NCTC10418. However, pKpQIL-D2 carriage in *S*. Typhimurium ATCC14028s conferred a significantly slower growth whereas the -UK variant did not. In contrast, the pKpQIL-UK plasmid significantly reduced growth rate of both *E. cloacae* and *S. marscescens* whereas the -D2 variant had no significant impact on the growth of these strains. These data show that there are differences in the fitness cost imposed by the two plasmid variants although this is species, and even strain specific suggesting a highly sensitive relationship between plasmid and host genotype and fitness.

Whilst impact on a host is important in determining the fitness of a plasmid so is the rate of transfer and number of strains which a plasmid can efficiently move into. Conjugation frequencies were determined for both variant plasmids. There were no major differences in frequency of transfer into *Salmonella* but the -UK variant was much more efficient in transferring into *E. coli* whereas the -D2 variant appeared to be more efficient in transferring into *Enterobacter* although this change did not reach statistical significance. These data may reflect changes in the impact on the host as well – the -UK variant imposed a significant fitness cost on Enterobacteriaceae, it may be that the -D2 variant is more efficient at entering these species, or this difference may be influenced by its relatively better fitness in these host species. The main processes of plasmid conjugation (DNA processing and horizontal transfer) are facilitated by genes encoded by the plasmid (31). Both the pKpQIL-UK and -D2 plasmids share identical transfer region and the only difference between the two plasmids is within the substituted region (17). We acknowledge that the donor clinical isolates used in this study were not isogenic. Hence, in addition to the substituted region in pKpQIL-D2, the differences in host background may also influence the transfer of the plasmids into the new recipient hosts observed in this study.

The imposition of fitness costs can result in a pressure to lose a plasmid, we also investigated stability of both variants in different hosts. Both plasmids were rapidly lost from *E. coli* populations but there were differences between the variant in other hosts. Although the plasmids were unable to persist within the *E. coli* NCTC10418 chosen in this study, this does not necessarily mean that pKpQIL-like plasmids will not be successful in other strains of this species. The pKpQIL-like plasmids have in fact been isolated from various *E. coli* clinical isolates (12, 13, 17). A previous study has also shown that a pKpQIL-like plasmid (pG12-KPC-2) introduced into uropathogenic *E. coli* ST10 (phylogroup A) persisted better than the ST69 (phylogroup D) counterpart (32). The loss of pKpQIL-D2 but not pKpQIL-UK from both *S*. Typhimurium hosts suggests that there is a fitness cost in *Salmonella* associated with carriage of the substituted region in pKpQIL-D2. The stability of large low copy plasmids within a bacterial population is generally facilitated by post-segregational killing and active partitioning mechanisms encoded by the plasmids (33). The pKpQIL-UK plasmid is known to encode genes involve in plasmid maintenance, such as the *parA*/*parB* and *stbA*/*stbB* (34). The variant plasmid, pKpQIL-D2 also carries the same *stbA*/*stbB* genes but a variant of *parA*/*parB* genes (17). It is possible that these genes were not able to function efficiently enough to maintain the plasmids in *Salmonella*. Although both plasmids had similar conjugation frequencies into the two *Salmonella* strains, pKpQIL-D2 was unable to persist. This may suggest that lateral gene transfer plays little role in the maintenance of these two plasmids. Taken together, our data and others (32, 35) reiterate that both plasmid and host factors play crucial roles in the persistence of a plasmid within a population.

In most host backgrounds there was no differential impact on antibiotic susceptibility from carriage of the variant plasmids, however the -UK variant conferred higher carbapenem MICs in *Serratia* than the -D2 variant. A previous study implicated the loss of OmpF and/or OmpC with the overproduction of AmpC β-lactamase as a cause for β-lactam resistance in *S. marcescens* (36). However, no difference in porin expression was observed in our study. Derepression of AmpC β-lactamase in the presence of antibiotic inducers has been previously reported. Some β-lactam antibiotics such as imipenem, cefoxitin and cefotetan are known to be strong AmpC β-lactamase inducers whilst third-generation cephalosporins such as cefotaxime and ceftazidime have poor inducing properties (37, 38). Therefore in *S. marcescens* NCTC10211 carrying the pKpQIL-D2 plasmid, the phenotype may be caused by incomplete derepression of the chromosomal AmpC β-lactamase.

Relative to the plasmid-free host bacterial strains, both plasmids affected the ability of various Enterobacteriaceae hosts to form a biofilm on plastic in different ways. Biofilm formation is an important phenomenon as most bacteria (99%) in the natural environment exist in biofilms (39). The establishment of a biofilm by a single or a mixture of bacterial species plays an important role in various infections and resistance to antimicrobial agents (antibiotics and biocides) (40). The true effect of plasmid carriage on the formation of biofilm is still unclear (41-43). Studies have suggested that conjugative factor(s) on plasmids may exert a global impact on the host affecting cell motility, cell-to-cell/surface interaction and curli production which are essential for the formation of mature biofilm (44, 45). In contrast, another study showed carriage of a conjugative plasmid pKJK5 caused a reduction in biofilm formation (43). In our study, carriage of either plasmid increased biofilm formation by *E. coli*, variable results were seen in *Salmonella* where the -UK variant improved biofilm formation by SL1344 but not 14028s; whilst, the -D2 variant had a negative impact on biofilm formed by 14028s. Again, this suggests differences between species and strains. The -D2 variant significantly increased the amount of biofilm formed by both *Serratia* and *Enterobacter*. Data from the crystal violet assay which is a measure of biomass and the flow cells where area coverage was measured did not always correlate – it may be denser biofilm forming strains did not spread as quickly.

Previously, we have carried out an in-depth transcriptomic profiling of *K. pneumoniae* ST258 carrying these two plasmids (19). The plasmids (in particular, the substituted region) affected the host transcriptomic landscape differently (19). Interestingly, the *K. pneumoniae* Ecl8 host carrying the pKpQIL-D2 plasmid out-competed the pKpQIL-UK counterpart in pairwise competition assay; even though the latter had a higher conjugation frequency than pKpQIL-D2 (19). Similar to the present study, we found no universal fitness benefit or cost imparted by the carriage of the pKpQIL-UK or -D2 plasmids across the collection of Enterobacteriaceae strains tested, highlighting that the fitness impact is a complex interaction between the plasmid and strain/species specific factors. A study competing an *E. coli* ST131 strain carrying an ESBL-encoding plasmid and its plasmid-cured variant with a commensal *E. coli* ST10 (IMT13353) found no difference in growth (46). However, the plasmid-cured variant of the *E. coli* ST131 was out-competed by another commensal *E. coli* ST10 (IMT13858) (46). This study (46) and others (30, 35) support our finding that the plasmid carriage impact on fitness to be multifactorial and difficult to extrapolate from one host-plasmid-environment relationship to another.

The widespread prevalence of *bla*_KPC_-carrying *K. pneumoniae* ST258-related clones poses a serious threat to medical treatment. At present, pKpQIL and its variants have played a significant role in the dissemination of this carbapenemase across various sequence type and species (12, 17). Although the reason for the successful spread and stability of this plasmid within a bacterial population still remains unclear, the findings of our study suggest that factors within the variable region of the plasmid can influence biologically important host phenotypes. It is plausible that antibiotic-independent selection pressure (47) coupled with the genetic fluidity of pKpQIL contributes to the effectiveness of this plasmid in spreading the *bla*_KPC_ gene. Our study provides better understanding of the interaction of plasmid factor(s) with different bacterial hosts background in determining the success of the plasmid. Although both plasmids studied here are very similar, the differences in the substituted region appear to confer biologically important differences in ability to transfer and persist within strains as well as influencing ability to form biofilms and in some species drug sensitivity. The emergence of variants of plasmids is well documented although the biological impact is not well understood. This study suggests that there are costs and benefits to plasmid alterations, which may provide access for a plasmid to new species but at a cost in others. Understanding how plasmids and hosts evolve requires much further study but is important if we want to be able to predict and, in the future, potentially manage the development of high fitness clones carrying important resistance genes on plasmids.

## MATERIALS AND METHODS

### Bacterial strains, plasmids, growth conditions and susceptibility to antibiotics

All plasmids and bacterial strains used in this study are listed in Table S2. Rifampicin-resistant mutants of *Escherichia coli* NCTC10418, *Salmonella* Typhimurium ATCC14028s and SL1344, *Enterobacter cloacae* NCTC10005 and *Serratia marcescens* NCTC10211 were selected as previously described (48) using 100 mg/L rifampicin. The plasmids were transferred into new host strains by filter-mating (49). All strains constructed were verified by PCR and DNA sequencing (Table S3). Throughout this study, ‘plasmid-free’ refers to the bacterial host before the introduction of pKpQIL-UK or -D2 plasmid. Unless otherwise stated, all experiments were carried out in LB (Miller) media.

The MICs of antibiotics for the strains used in this study were determined using the guidelines recommended by the British Society for Antimicrobial Chemotherapy (BSAC) (50). *E. coli* NCTC10418 was used as the control strain. A difference in the MIC of antibiotics of more than one 2-fold dilution for strains with and without a plasmid was considered significant.

### Growth kinetics

Bacterial growth during the logarithmic phase was used to determine the fitness of the host strain when a plasmid had been introduced into the strain (49). Briefly, an overnight culture was diluted with fresh broth to give a final inoculum of 4% (v/v) and 200 µl was transferred into the wells of a 96-well microtitre plate (Sterilin). The growth of the cultures was monitored at OD600 at 10 min intervals per cycle for 100 cycles using the FLUOstar Optima plate reader (BMG Labtech). The generation times were determined on three separate occasions. The value for the plasmid-free strain was compared with that for the plasmid-containing strain using the Student’s *t*-test; *p* values less than 0.05 were considered statistically significant.

### Determination of conjugation frequency

The conjugation frequency of plasmids from their original clinical isolates was determined as previously described (49). Cultures of the donor and recipient bacterial strains were grown at 37°C with shaking (200 rpm) until they had reached mid-logarithmic phase. Cultures were standardised to an OD600 of 0.6 and mixed together in a total volume of 100 μl at a 1:2 donor-to-recipient ratio. The suspension was then transferred on to sterile 0.45 μm nylon filters (Millipore) and placed on the surface of an agar plate and incubated at 37°C for 3 hours. The bacteria were removed from the filters by vigorous agitation in 1 mL of LB medium, and serially diluted and plated on agar supplemented with selecting antibiotics (0.25 mg/L doripenem and 100 mg/L rifampicin) and incubated overnight at 37°C. The conjugation frequency of the plasmids was expressed as the number of transconjugants (cfu/mL) over half of the number of donors (cfu/mL).

The frequency of conjugation was calculated on three separate occasions and differences between donor strains or different plasmids for the same host strain were deemed significant when p<0.05 by Student’s *t*-test.

### Plasmid persistence

The proportion of the bacterial population that retained the plasmid was determined over a period of 20 days as previously described (49). A single colony was inoculated in 10 mL broth supplemented with 0.25 mg/L doripenem and incubated overnight at 37°C at 200 rpm. The culture was sub-cultured daily into fresh broth without antibiotics at a dilution of 1 in 100 over a period of 20 days. At timed intervals (day 5, 10, 15 and 20), the culture was serially diluted and plated on agar. The following day, colonies were replica-plated on to agar supplemented with 0.25 mg/L doripenem. The retention of the plasmid in a particular bacterial strain was calculated as a percentage of colonies on doripenem agar over the total number of colonies observed on the antibiotic-free LB agar. The experiment was repeated on three separate occasions.

### Biofilm formation on plastic

The impact of plasmid carriage on biofilm formation by the host bacterial strains was assessed in 96-well microtitre trays as previously described (51). Strains were assessed for biofilm formation in LB broth without sodium chloride, which allows efficient biofilm formation by Enterobacteriaceae (52). Diluted overnight culture (200 μl) at a final inoculum of 4% (v/v) was transferred into the wells of the microtitre tray and incubated at 30°C with gentle shaking for 48 hours. The biofilm formed in the wells was stained with 0.1% (w/v) crystal violet solution (Sigma Aldrich) for 15 minutes and washed with sterile water to remove the excess dye. The crystal violet stain was solubilised in 200 μl of 70% (v/v) ethanol with gentle shaking for 15 min. The intensity of the solubilised crystal violet, which relates to the amount of biofilm formed, was measured at OD600. The ability of each strain, with and without a plasmid, to form a biofilm was determined in three independent experiments. Data were analysed using the Student’s *t*-test and differences between strains with and without a plasmid of p<0.05 were considered as significant.

### Biofilm formation under constant flow

The ability of the plasmid-carrying bacteria to form biofilms was also assessed under constant flow of liquid media in glass microfluidic flow cells as previously described (51) with some modifications. Overnight cultures of the strains were diluted to an OD600 of 0.1. To allow the bacterial strains to attach to the surface of the microfluidic channel, after inoculation the 48-well plate was incubated on a heating block at 30°C for 2 hours. After incubation, 1 mL of fresh broth (without sodium chloride) was added into the input well and the flow rate was set to a constant 0.3 dynes/min and the plate was incubated on the heating block for 48 hours. The biofilms were observed at 6, 12, 24 and 48-hour time points at 40X magnification. The area of coverage by the biofilm in the microfluidic channel was determined using image analysis software, ImageJ (http://imagej.nih.gov/ij/). The Student’s *t-*test was used to determine whether a significant difference (p<0.05) was observed between strains with and without plasmids in the area of coverage.

## Supporting information

## ACKNOWLEDGEMENTS

This project was supported by the Elite Doctoral Researcher Scholarship of the University of Birmingham awarded to L.J.V.P to support H.T.H.S. H.T.H.S performed the experiments.

H.T.H.S, M.A.W & L.J.V.P designed the experiments and analysed the data. All authors contributed to writing the manuscript. L.J.V.P conceived and supervised the project.

H.T.H.S, M.A.W and L.J.V.P have none to declare. N.W has no personal conflicts of interest to declare. However, PHE’s AMRHAI Reference Unit has received financial support for conference attendance, lectures, research projects or contracted evaluations from numerous sources, including: Accelerate Diagnostics, Achaogen Inc., Allecra Therapeutics, Amplex, AstraZeneca UK Ltd, AusDiagnostics, Basilea Pharmaceutica, Becton Dickinson Diagnostics, bioMérieux, Bio-Rad Laboratories, The BSAC, Cepheid, Check-Points B.V., Cubist Pharmaceuticals, Department of Health, Enigma Diagnostics, European Centre for Disease Prevention and Control, Food Standards Agency, GlaxoSmithKline Services Ltd, Helperby Therapeutics, Henry Stewart Talks, IHMA Ltd, Innovate UK, Kalidex Pharmaceuticals, Melinta Therapeutics, Merck Sharpe & Dohme Corp, Meiji Seika Pharma Co., Ltd, Mobidiag, Momentum Biosciences Ltd, Neem Biotech, NIHR, Nordic Pharma Ltd, Norgine Pharmaceuticals, Rempex Pharmaceuticals Ltd, Roche, Rokitan Ltd, Smith & Nephew UK Ltd, Shionogi & Co. Ltd, Trius Therapeutics, VenatoRx Pharmaceuticals, Wockhardt Ltd., and the World Health Organization.

## Supplementary Material

**Table S1.**
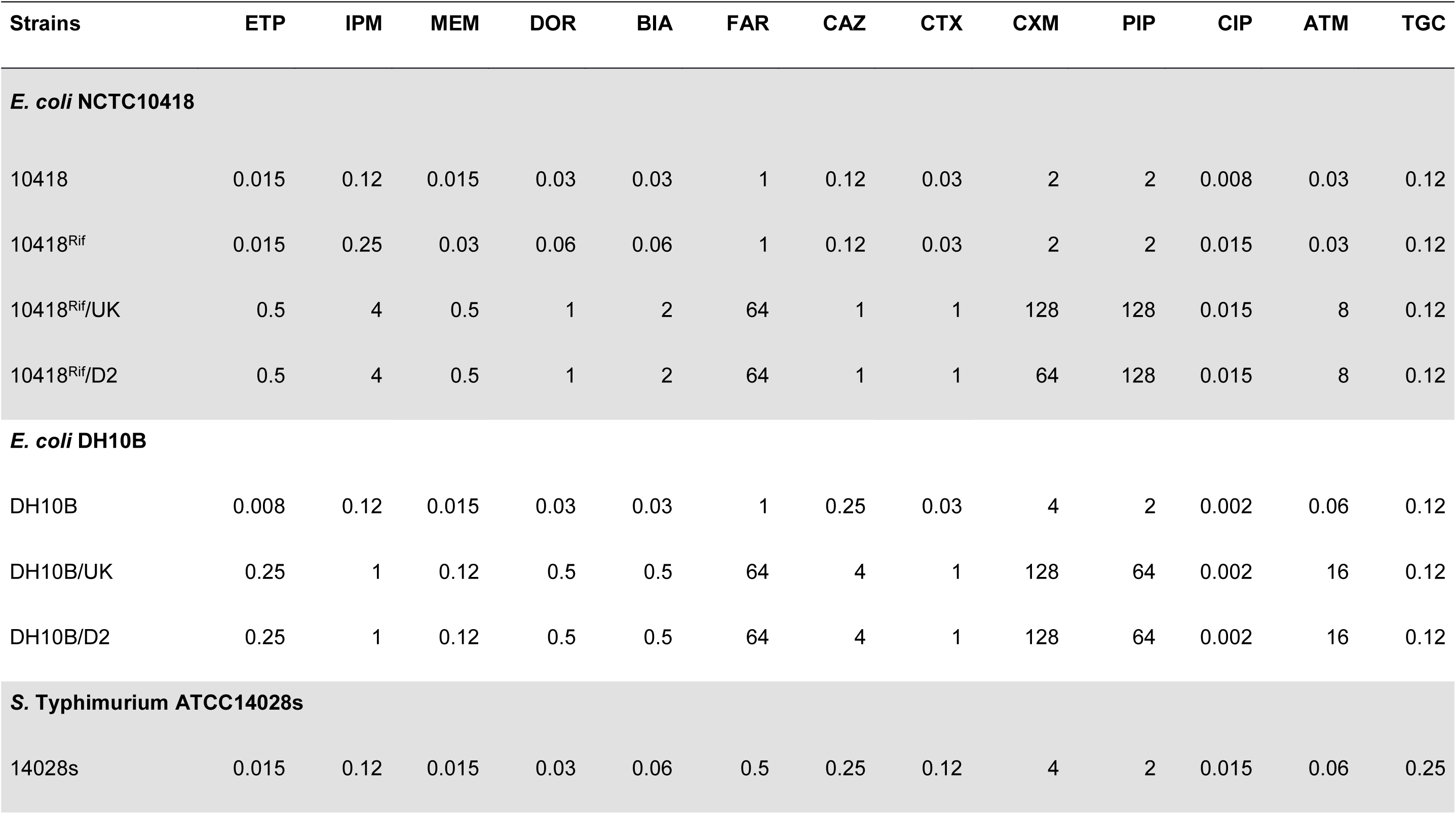
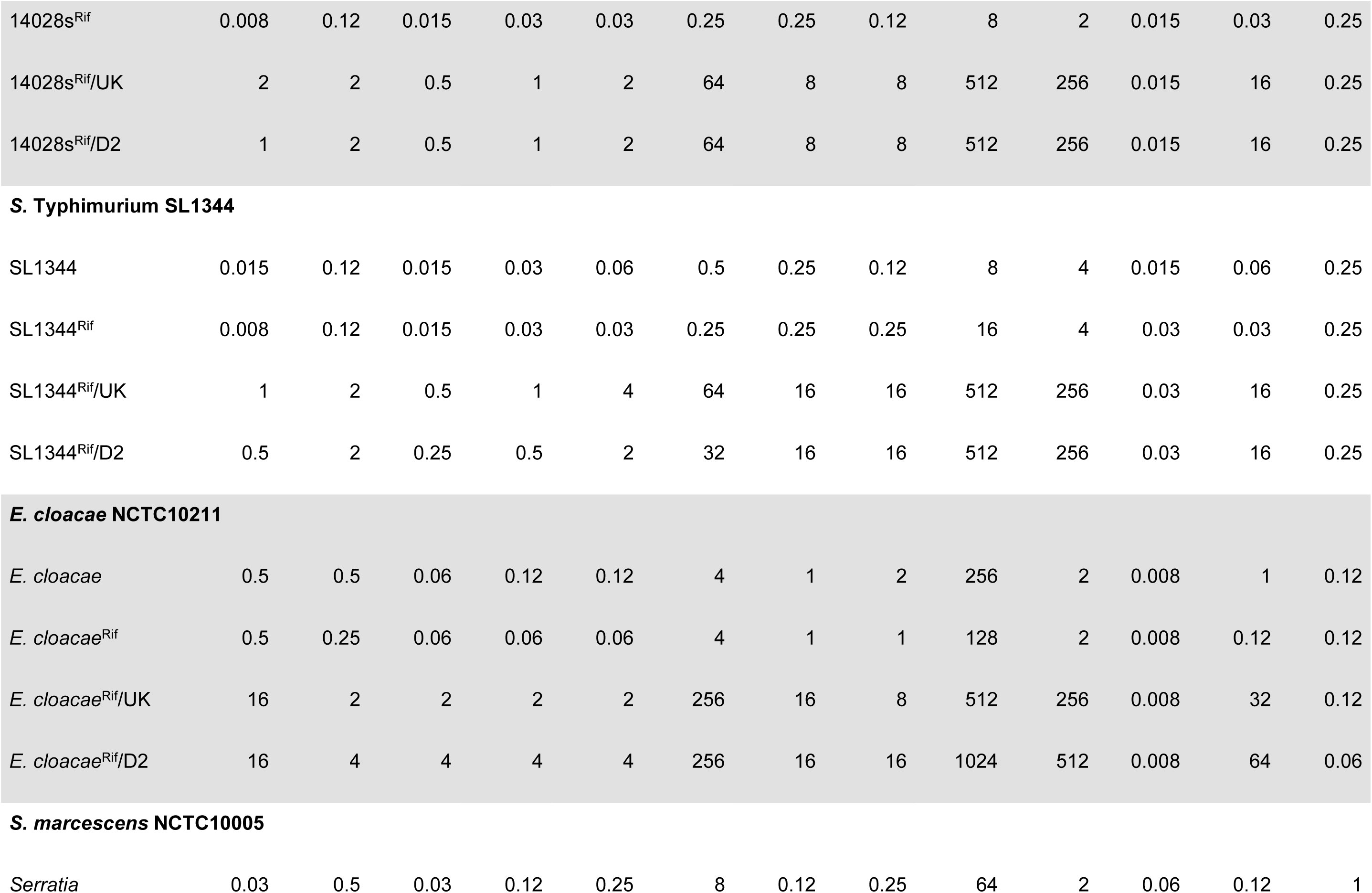
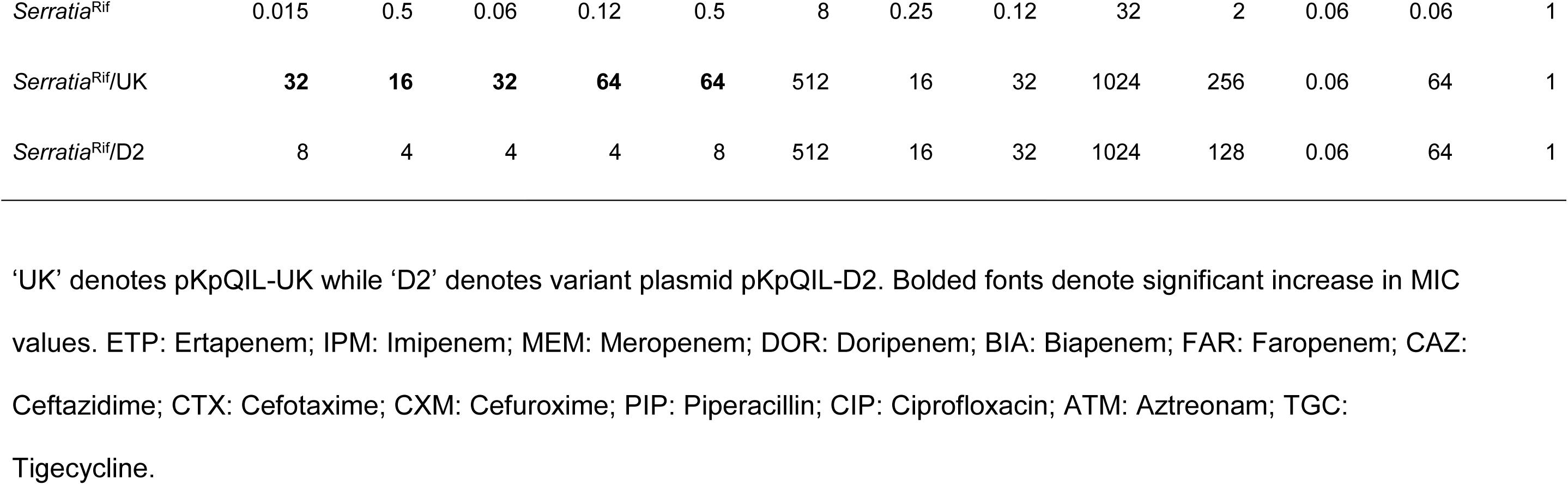
MICs (mg/L) of antibiotics for hosts carrying pKpQIL-UK and -D2.

**Table S2.**
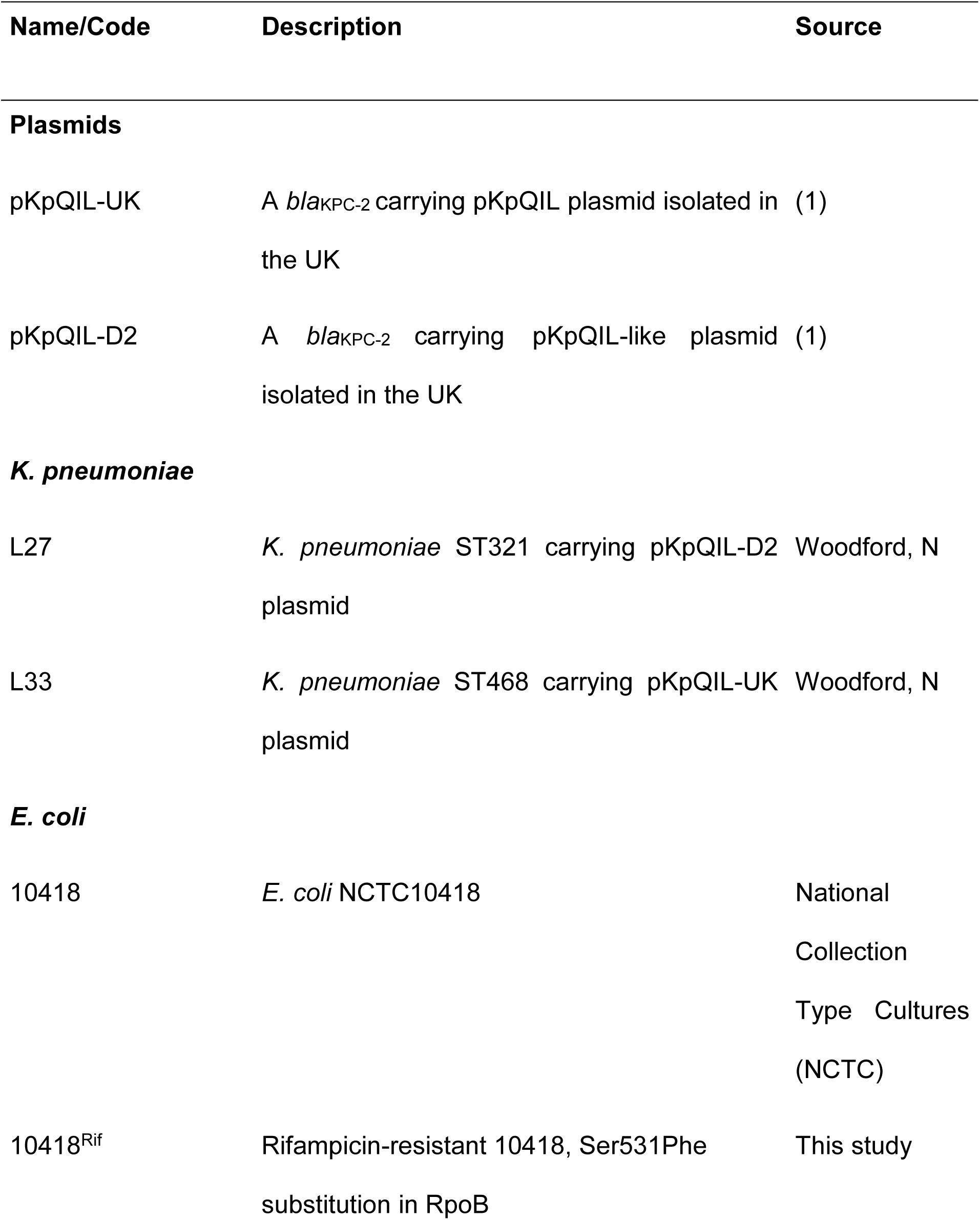
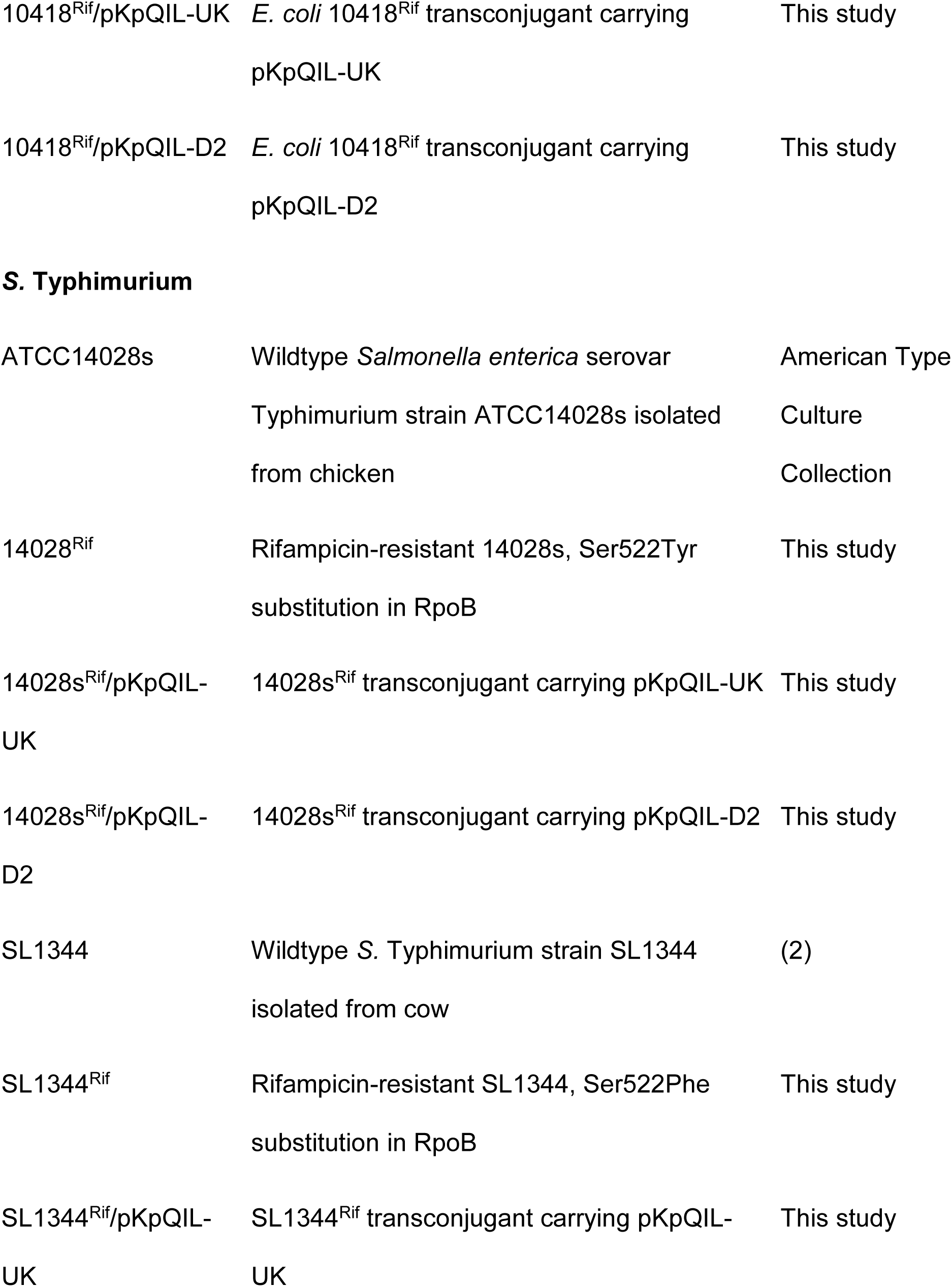
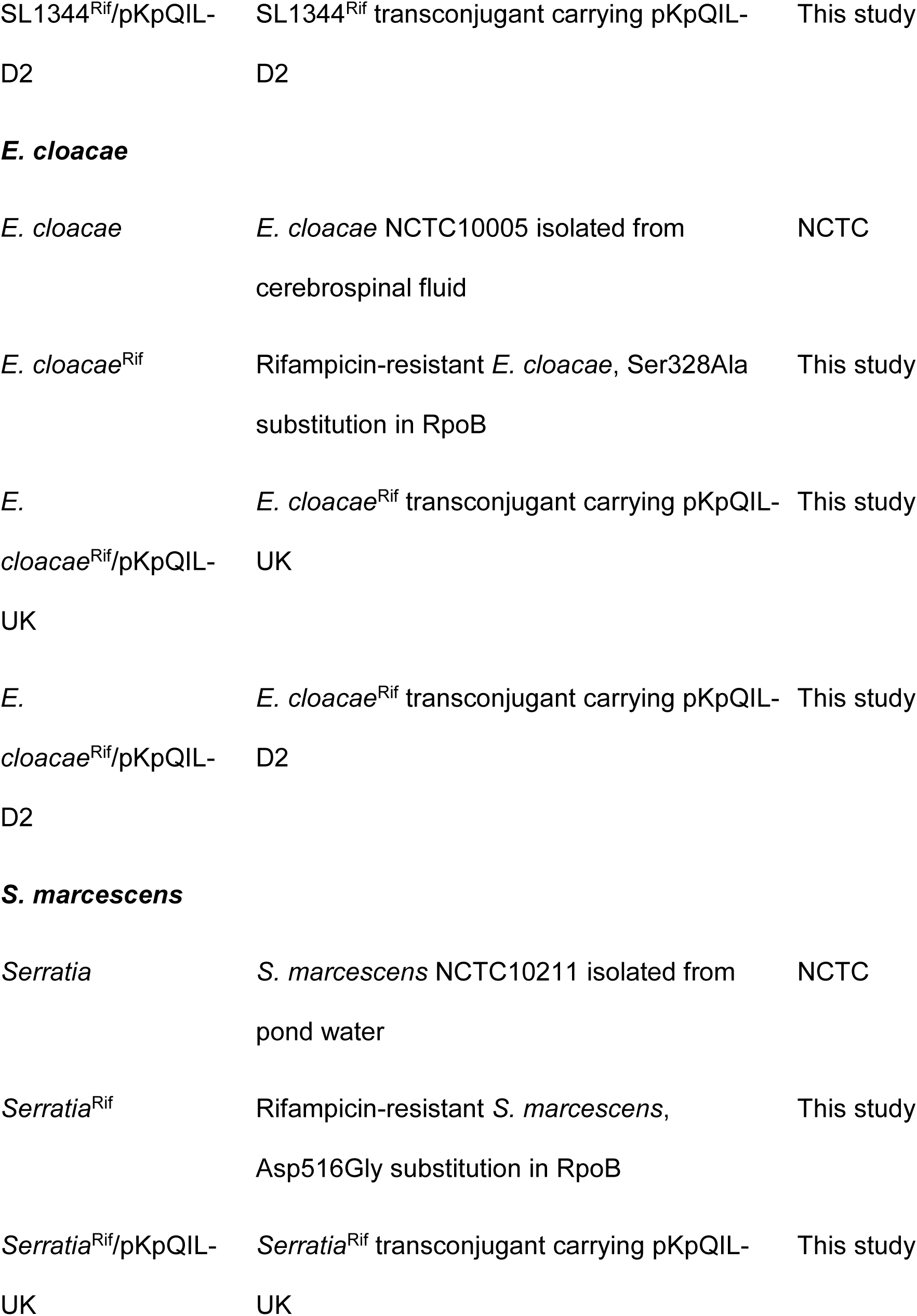
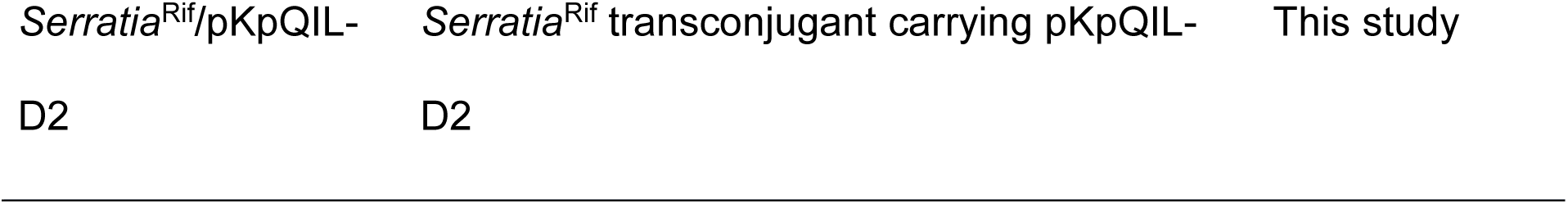
Bacterial strains and plasmids used in this study.

**Table S3.**
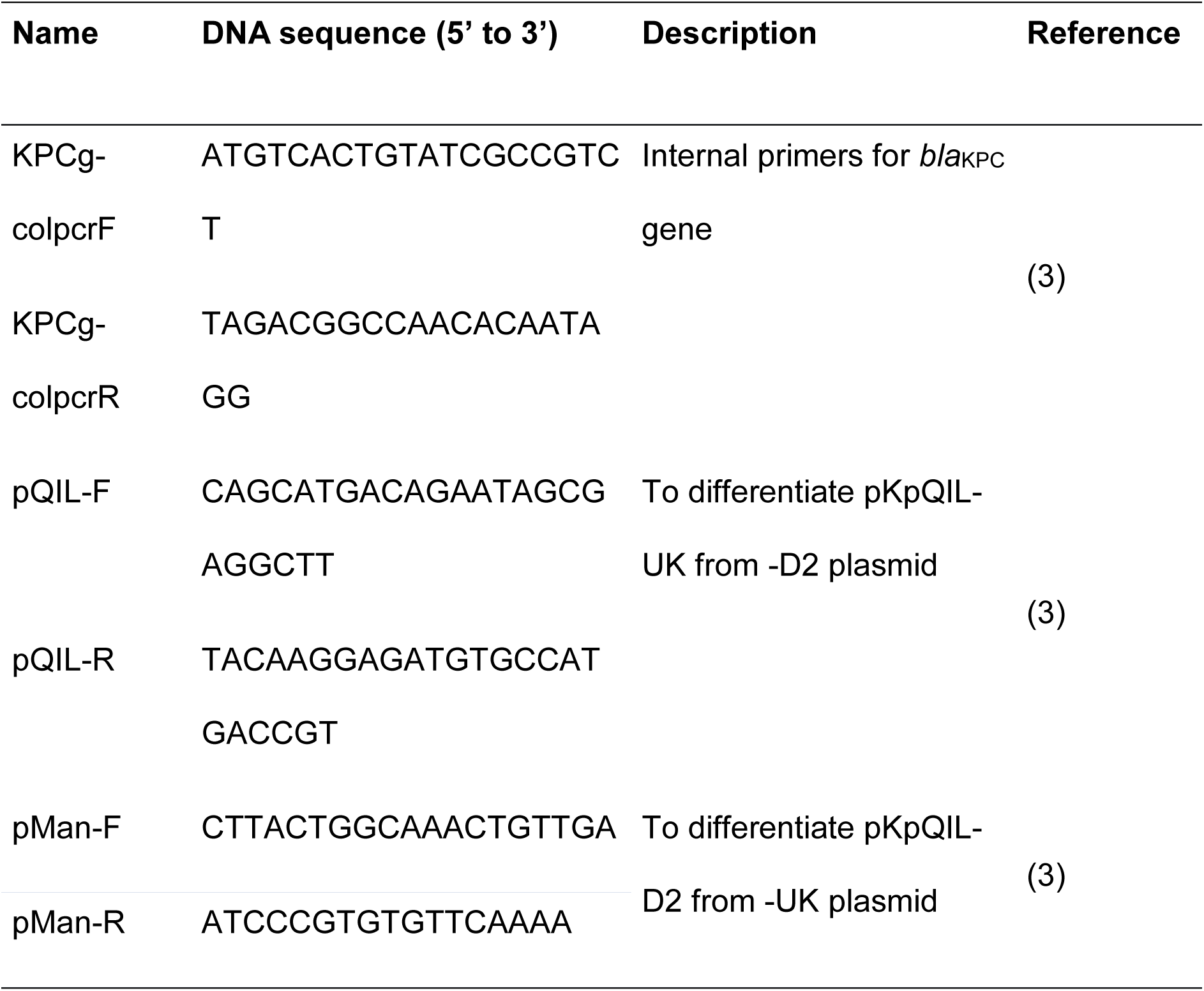
List of primers used in this study.

