## Supplementary material for "Different impacts of naturally occurring variants of the globally disseminated carbapenemase-encoding plasmid pKpQIL on the biology of host strains"

### Table S1. MICs (mg/L) of antibiotics for hosts carrying pKpQIL-UK and -D2

| Strains | ETP | IPM | MEM | DOR | BIA | FAR | CAZ | CTX | CXM | PIP | CIP | ATM | TGC |
|---|---|---|---|---|---|---|---|---|---|---|---|---|---|
| ***E. coli*** **NCTC10418** | | | | | | | | | | | | | |
| 10418 | 0.015 | 0.12 | 0.015 | 0.03 | 0.03 | 1 | 0.12 | 0.03 | 2 | 2 | 0.008 | 0.03 | 0.12 |
| 10418$^{Rif}$ | 0.015 | 0.25 | 0.03 | 0.06 | 0.06 | 1 | 0.12 | 0.03 | 2 | 2 | 0.015 | 0.03 | 0.12 |
| 10418$^{Rif}$/UK | 0.5 | 4 | 0.5 | 1 | 2 | 64 | 1 | 1 | 128 | 128 | 0.015 | 8 | 0.12 |
| 10418$^{Rif}$/D2 | 0.5 | 4 | 0.5 | 1 | 2 | 64 | 1 | 1 | 64 | 128 | 0.015 | 8 | 0.12 |
| ***E. coli*** **DH10B** | | | | | | | | | | | | | |
| DH10B | 0.008 | 0.12 | 0.015 | 0.03 | 0.03 | 1 | 0.25 | 0.03 | 4 | 2 | 0.002 | 0.06 | 0.12 |
| DH10B/UK | 0.25 | 1 | 0.12 | 0.5 | 0.5 | 64 | 4 | 1 | 128 | 64 | 0.002 | 16 | 0.12 |
| DH10B/D2 | 0.25 | 1 | 0.12 | 0.5 | 0.5 | 64 | 4 | 1 | 128 | 64 | 0.002 | 16 | 0.12 |
| ***S.*** **Typhimurium ATCC14028s** | | | | | | | | | | | | | |
| 14028s | 0.015 | 0.12 | 0.015 | 0.03 | 0.06 | 0.5 | 0.25 | 0.12 | 4 | 2 | 0.015 | 0.06 | 0.25 |
| 14028s$^{Rif}$ | 0.008 | 0.12 | 0.015 | 0.03 | 0.03 | 0.25 | 0.25 | 0.12 | 8 | 2 | 0.015 | 0.03 | 0.25 |
| 14028s$^{Rif}$/UK | 2 | 2 | 0.5 | 1 | 2 | 64 | 8 | 8 | 512 | 256 | 0.015 | 16 | 0.25 |
| 14028s$^{Rif}$/D2 | 1 | 2 | 0.5 | 1 | 2 | 64 | 8 | 8 | 512 | 256 | 0.015 | 16 | 0.25 |
| ***S.*** **Typhimurium SL1344** | | | | | | | | | | | | | |
| SL1344 | 0.015 | 0.12 | 0.015 | 0.03 | 0.06 | 0.5 | 0.25 | 0.12 | 8 | 4 | 0.015 | 0.06 | 0.25 |
| SL1344$^{Rif}$ | 0.008 | 0.12 | 0.015 | 0.03 | 0.03 | 0.25 | 0.25 | 0.25 | 16 | 4 | 0.03 | 0.03 | 0.25 |
| SL1344$^{Rif}$/UK | 1 | 2 | 0.5 | 1 | 4 | 64 | 16 | 16 | 512 | 256 | 0.03 | 16 | 0.25 |
| SL1344$^{Rif}$/D2 | 0.5 | 2 | 0.25 | 0.5 | 2 | 32 | 16 | 16 | 512 | 256 | 0.03 | 16 | 0.25 |

| | | | | | | | | | | | | | |
|---|---|---|---|---|---|---|---|---|---|---|---|---|---|
| ***E. cloacae* NCTC10211** | | | | | | | | | | | | | |
| *E. cloacae* | 0.5 | 0.5 | 0.06 | 0.12 | 0.12 | 4 | 1 | 2 | 256 | 2 | 0.008 | 1 | 0.12 |
| *E. cloacae*$^{Rif}$ | 0.5 | 0.25 | 0.06 | 0.06 | 0.06 | 4 | 1 | 1 | 128 | 2 | 0.008 | 0.12 | 0.12 |
| *E. cloacae*$^{Rif}$/UK | 16 | 2 | 2 | 2 | 2 | 256 | 16 | 8 | 512 | 256 | 0.008 | 32 | 0.12 |
| *E. cloacae*$^{Rif}$/D2 | 16 | 4 | 4 | 4 | 4 | 256 | 16 | 16 | 1024 | 512 | 0.008 | 64 | 0.06 |
| ***S. marcescens* NCTC10005** | | | | | | | | | | | | | |
| *Serratia* | 0.03 | 0.5 | 0.03 | 0.12 | 0.25 | 8 | 0.12 | 0.25 | 64 | 2 | 0.06 | 0.12 | 1 |
| *Serratia*$^{Rif}$ | 0.015 | 0.5 | 0.06 | 0.12 | 0.5 | 8 | 0.25 | 0.12 | 32 | 2 | 0.06 | 0.06 | 1 |
| *Serratia*$^{Rif}$/UK | **32** | **16** | **32** | **64** | **64** | 512 | 16 | 32 | 1024 | 256 | 0.06 | 64 | 1 |
| *Serratia*$^{Rif}$/D2 | 8 | 4 | 4 | 4 | 8 | 512 | 16 | 32 | 1024 | 128 | 0.06 | 64 | 1 |

‘UK’ denotes pKpQIL-UK while ‘D2’ denotes variant plasmid pKpQIL-D2. Bolded fonts denote significant increase in MIC values. ETP: Ertapenem; IPM: Imipenem; MEM: Meropenem; DOR: Doripenem; BIA: Biapenem; FAR: Faropenem; CAZ: Ceftazidime; CTX: Cefotaxime; CXM: Cefuroxime; PIP: Piperacillin; CIP: Ciprofloxacin; ATM: Aztreonam; TGC: Tigecycline.

**Table S2. Bacterial strains and plasmids used in this study.**

| Name/Code | Description | Source |
|---|---|---|
| **Plasmids** | | |
| pKpQIL-UK | A *bla*$_{KPC-2}$ carrying pKpQIL plasmid isolated in the UK | (1) |
| pKpQIL-D2 | A *bla*$_{KPC-2}$ carrying pKpQIL-like plasmid isolated in the UK | (1) |
| ***K. pneumoniae*** | | |
| L27 | *K. pneumoniae* ST321 carrying pKpQIL-D2 plasmid | Woodford, N |
| L33 | *K. pneumoniae* ST468 carrying pKpQIL-UK plasmid | Woodford, N |
| ***E. coli*** | | |
| 10418 | *E. coli* NCTC10418 | National Collection Type Cultures (NCTC) |
| 10418$^{Rif}$ | Rifampicin-resistant 10418, Ser531Phe substitution in RpoB | This study |
| 10418$^{Rif}$/pKpQIL-UK | *E. coli* 10418$^{Rif}$ transconjugant carrying pKpQIL-UK | This study |
| 10418$^{Rif}$/pKpQIL-D2 | *E. coli* 10418$^{Rif}$ transconjugant carrying pKpQIL-D2 | This study |
| ***S.* Typhimurium** | | |
| ATCC14028s | Wildtype *Salmonella enterica* serovar Typhimurium strain ATCC14028s isolated from chicken | American Type Culture Collection |
| 14028$^{Rif}$ | Rifampicin-resistant 14028s, Ser522Tyr substitution in RpoB | This study |

| | | |
|---|---|---|
| 14028s$^{Rif}$/pKpQIL-UK | 14028s$^{Rif}$ transconjugant carrying pKpQIL-UK | This study |
| 14028s$^{Rif}$/pKpQIL-D2 | 14028s$^{Rif}$ transconjugant carrying pKpQIL-D2 | This study |
| SL1344 | Wildtype *S.* Typhimurium strain SL1344 isolated from cow | (2) |
| SL1344$^{Rif}$ | Rifampicin-resistant SL1344, Ser522Phe substitution in RpoB | This study |
| SL1344$^{Rif}$/pKpQIL-UK | SL1344$^{Rif}$ transconjugant carrying pKpQIL-UK | This study |
| SL1344$^{Rif}$/pKpQIL-D2 | SL1344$^{Rif}$ transconjugant carrying pKpQIL-D2 | This study |
| ***E. cloacae*** | | |
| *E. cloacae* | *E. cloacae* NCTC10005 isolated from cerebrospinal fluid | NCTC |
| *E. cloacae*$^{Rif}$ | Rifampicin-resistant *E. cloacae*, Ser328Ala substitution in RpoB | This study |
| *E. cloacae*$^{Rif}$/pKpQIL-UK | *E. cloacae*$^{Rif}$ transconjugant carrying pKpQIL-UK | This study |
| *E. cloacae*$^{Rif}$/pKpQIL-D2 | *E. cloacae*$^{Rif}$ transconjugant carrying pKpQIL-D2 | This study |
| ***S. marcescens*** | | |
| *Serratia* | *S. marcescens* NCTC10211 isolated from pond water | NCTC |
| *Serratia*$^{Rif}$ | Rifampicin-resistant *S. marcescens*, Asp516Gly substitution in RpoB | This study |
| *Serratia*$^{Rif}$/pKpQIL-UK | *Serratia*$^{Rif}$ transconjugant carrying pKpQIL-UK | This study |
| *Serratia*$^{Rif}$/pKpQIL-D2 | *Serratia*$^{Rif}$ transconjugant carrying pKpQIL-D2 | This study |

**Table S3. List of primers used in this study.**

| Name | DNA sequence (5' to 3') | Description | Reference |
|---|---|---|---|
| KPCg-colpcrF | ATGTCACTGTATCGCCGTCT | Internal primers for $bla_{KPC}$ gene | (3) |
| KPCg-colpcrR | TAGACGGCCAACACAATAGG | | |
| pQIL-F | CAGCATGACAGAATAGCGAGGCTT | To differentiate pKpQIL-UK from -D2 plasmid | (3) |
| pQIL-R | TACAAGGAGATGTGCCATGACCGT | | |
| pMan-F | CTTACTGGCAAACTGTTGA | To differentiate pKpQIL-D2 from -UK plasmid | (3) |
| pMan-R | ATCCCGTGTGTTCAAAA | | |
